# A Comprehensive Workflow for Imaging Live Insulin Secretion Events and Granules in Intact Islets

**DOI:** 10.1101/2025.04.22.650066

**Authors:** Margret A. Fye, Rahul Sharma, Phyu Sin M. Myat, Pi’ilani Regan, Hudson McKinney, Guoqiang Gu, Irina Kaverina

## Abstract

Accurate detection of insulin secretion from pancreatic beta cells is crucial for understanding normal physiological insulin secretion and its pathophysiological counterpart in diabetic states. Traditional methods using fluorescently labeled insulin granules or dye labeling often struggle to distinguish secretion from insulin granule dynamics. We present an optimized protocol using the cell-impermeable Zn^2+^-binding dye FluoZin-3, which fluoresces upon Zn^2+^ co-secretion with insulin outside of the islet, more accurately representing secretion. FluoZin-3 combined with intact islet attachment to vascular extracellular matrix and TIRF microscopy offers high spatial and temporal resolution as well as a high signal-to-noise ratio in a minimally perturbed system. Additionally, by integrating the cell-permeable Zn^2+^-binding dye ZIGIR, we can track insulin granule dynamics alongside secretion events. Our approach generates large datasets, which we efficiently analyze using ilastik machine learning software, enabling fast, accurate, and optionally supervised analysis. This technique builds on our group’s previous protocols, detailing a streamlined workflow adaptable to high-resolution, live-cell microscopy for not just insulin but other secretory/granule systems as well. With this method, we investigated secretion behavior of different IG pools in real time during the first phase of insulin secretion: predocked, which appear before high glucose stimulation and are docked at the membrane; docked, which appear upon high glucose stimulation and dock at the membrane; and newcomer, which appear upon high glucose stimulation but don’t dock at the membrane. The predocked and newcomer insulin granules are equally secreted and newcomer insulin granules dwell less than one second before secretion upon high glucose stimulation. The predocked and docked insulin granules, however, stay longer at the membrane before secretion. This method is useful for the investigation of functional beta cell heterogeneity of insulin granule secretion in space and time.

## Introduction

Insulin secretion from pancreatic beta cells is a critical process for normal glucose homeostasis and therefore accuracy in capturing insulin secretion events for study is of the utmost importance. Live imaging of fluorescently-labeled insulin granules (IGs) allows for localization of granules while inside the beta cell in real-time^1–3^, but this technique is limited in its ability to visualize secretion events, relying on disappearance of IGs which is equally likely due to withdrawal of IGs from docking sites. Other studies employ label-free approaches^4, 5^ or use dye-based approaches such as sulforhodamine B to label the exocytic cavity^6, 7^. However, using rhodamine-based dye uptake introduces a similar challenge, that it becomes difficult to separate exocytosis from endocytosis. These issues can be circumvented by using a cell-impermeable dye, notably the FluoZin-3 dye^8^, which binds Zn^2+^ co-secreted with insulin and subsequently fluoresces green, giving off bright, transient signal that is simple to localize at a sub-pixel level. Our group has optimized this label-free approach to record IG exocytosis with high spatial (sub-pixel) and temporal (millisecond) resolution^9, 10^. This method is advantageous due to its lack of interference with insulin itself, clear exocytosis labeling, and simple microscopy technique. Moreover, insulin secretion is preferentially directed toward the vasculature^7, 11^, and attaching islets to vascular extracellular matrix (ECM) preserves this phenomenon while also promoting beta cell fitness in intact, isolated islets^12, 13^. Our method using total internal reflection fluorescence (TIRF) microscopy allows for optimal visualization of this beta cell-vascular interface at a high signal-to-noise ratio, while reducing photobleaching and toxicity in the islets.

A drawback of this approach, though, is that it precludes the use of any green fluorescent insulin fusion proteins to assess IG dynamics prior to secretion. However, we have here combined our previously described technique with the cell-permeable, Zn^2+^-specific dye ZIGIR^14, 15^ which was more recently published and made available. This provides us a significant edge in analyzing IG dynamics relative to secretion events which was previously challenging due to lack of existing materials.

This technique, while highly efficacious, faces the issue of generating large and complex datasets which are time-consuming to process and analyze. In our previous publications these datasets were manually analyzed^9, 10^, but with recent advances in machine learning technology, we have been able to analyze these datasets quickly and accurately, while maintaining user supervision^16^, using the machine learning freeware ilastik^17^. This machine learning software was adaptable into our existing protocols and MATLAB script for analysis, making it highly effective for analyzing FluoZin-3-generated datasets. Here we describe a continuation of the islet isolation and attachment protocol described in Ho et al. (2023)^18^, in which we detail the FluoZin-3 and ZIGIR protocol itself, consisting of pre-Ilastik processing, ZIGIR IG dynamics processing, Ilastik training and batch processing, and Ilastik information export for input into our clustering MATLAB script. This workflow is highly adaptable and applicable to any high-resolution live-cell microscopy involving Zn^2+^ secretion, automated fluorescence transient signal detection, and clustering. Using this methodology, we here investigate secretion heterogeneity of different IG pools (predocked, docked, and newcomer) in beta cells.

### Development of the protocol

Towards developing a better method for assessing and imaging live secretion events, our group drew upon the methods originally published in Gee et al. (2002)^8^. Capitalizing on the knowledge that Zn^2+^ ions co-crystallize with insulin peptide, this publication developed a Zn^2+^ indicator that shared molecular moieties with the known Ca^2+^ sensors fluo-3 and fluo-4 but enhanced its affinity for Zn^2+^ substantially over Ca^2+^, conferring its specificity. Working as a weak chelator, this probe has excitation/emission peaks of 488/520 nm, is cell impermeable, and is virtually non-fluorescent in the absence of Zn^2+^ with a several-hundred-fold increase in fluorescence upon Zn^2+^ introduction to solution^8^. These characteristics make it a perfect candidate for assessing insulin secretion from beta cells, whereupon Zn^2+^ is released extracellularly when insulin crystals dissolve in solution.

Using this as a basis for developing our protocol and the precedent for attaching to vascular ECM^19^, Xhu et al. (2015)^9^ describes our group’s first use of this technique. Adding on to this protocol, our group used intact mouse islets rather than the originally proposed isolated beta cells, to assess secretion in an *in situ* environment. We further used Fiji software to identify flashes in time and space and obtain precise coordinates of these flashes that facilitated analysis of insulin secretion following modulations to the microtubule cytoskeleton^9^.

After observations that several of these events clustered in time and space, our group next adapted this protocol to include a clustering analysis in Trogden et al. (2021)^10^. This analysis includes the original time and space insulin secretion quantification, with the addition of a custom MATLAB script that imports these coordinates and using a density-based scanning (DBS) algorithm with nearest neighbor analysis. The DBS algorithm includes parameters to identify a “cluster” of insulin secretion events based on a minimum of three neighbors within an event, with a distance radius of 1.5µm. This allowed us to identify clusters of secretion events that we termed “hot spots,” a phenomenon that has been since replicated in other studies and has allowed for complex analysis of the heterogeneity of single beta cells within an islet.

Further studies sought to expedite this analysis for high data volumes, as the protocol in Xhu and Trogden involved manual identification of flashes. In Fye et al. (2025)^16^ we trained the machine learning image analysis software ilastik to automatically identify flashes using sequential pixel then object classification workflows. This streamlining of the process allowed for rapid, accurate, and blind identification of insulin secretion events to improve the overall method.

The final addition to this workflow was the ability to track insulin granule movement prior to secretion. We have thus far been limited by the FluoZin-3 assay’s ability to only label extracellular events, while other insulin probes are limited by their ability to only label granules within the cell. We sought to bridge this gap by using another Zn^2+^-based dye, ZIGIR. ZIGIR is cell-permeable and binds Zn^2+^ within dense core granules, allowing visualization intracellularly^14,15^. Additionally, ZIGIR is a red dye as compared to FluoZin-3 being a green dye, making it optimal for dual imaging where in comparison other green dyes are not often compatible with FluoZin-3 as it creates too much background signal. We have recently developed a protocol in which intact islets are pre-treated with ZIGIR to label insulin granules, and are subsequently stimulated with glucose and FluoZin-3 to visualize both granule motility and secretion concurrently. From this, we have been able to observe the movement of a granule prior to its secretion.

### Applications of the method

Although this method was originally developed for studying beta cell insulin secretion, it has potential for wider applications. As a valuable method for imaging both intracellular and extracellular Zn^2+^, it opens up possibilities for imaging any system that may have significant concentrations of Zn^2+^, with particular focus on secretory heterogeneity and docking/secretion behavior. Additionally, our method for automatic detection of secretion events could be applicable to other secretory imaging based assays, such as neurotransmitter secretion from neurons. As ilastik allows high adaptability, flexibility, and trainability, this particular aspect of the workflow is optimal for any secretion system. This workflow is not only effective for intact islet systems, but is adaptable to other single cell or tissue systems.

### Comparison with other methods

Some other common methods for detecting insulin secretion events include use of insulin granule probes followed by insulin granule disappearance (ratiometric or degranulation) or extracellular dye uptake into the exocytic cavity.

Several probes for insulin granules exist, though each present unique benefits and drawbacks. Insulin is a relatively large molecule and gets packed tightly into granules after synthesis, making it difficult for proper insulin folding and granule maturation^20^. Some common markers of IGs are C-peptide, a processing result of insulin maturation^21^; phogrin^22, 23^; and neuropeptide Y^24, 25^, among others. While these markers can faithfully label insulin granules within live beta cells, the drawback to using these is that they are either unable to faithfully show where secretion occurs and/or show granule disappearance. There are also examples of overexpression of these proteins impairing the normal GSIS pathway, as in the case of phogrin^23^. In particular, identifying granule disappearance as secretion can be erroneous as evidenced by our group’s finding that granule disappearance is often a result of IG trafficking away from sites of secretion, along MTs^9, 26, 27^. In this case, not all events can be ascribed to secretion, and labeling these events as secretion would skew the data. Our method, on the other hand, labels these events outside of the cell, concurrent with their occurrence making it significantly more reliable for detecting actual secretion events rather than disappearance of intracellular granules.

The use of fluorescent dyes to label insulin secretion is often favorable as it interferes less with the normal GSIS pathway. A common method is to use the cell impermeable dye sulforhodamine B which will increase in fluorescence upon opening of the exocytic pore and its entrance therein. Signal will then be quenched upon completion of secretion and fusion of the granule with the membrane^6, 7^. Although this is a useful method, it presents difficulty in distinguishing exocytosis and endocytosis. On the other hand, labeling extracellularly released Zn^2+^ can only be associated with secretion, as endocytosis would not show a similar release.

### Experimental Design

Here we outline a comprehensive workflow for assessing insulin secretion from intact islets (Figure 1).

**Figure 1.**
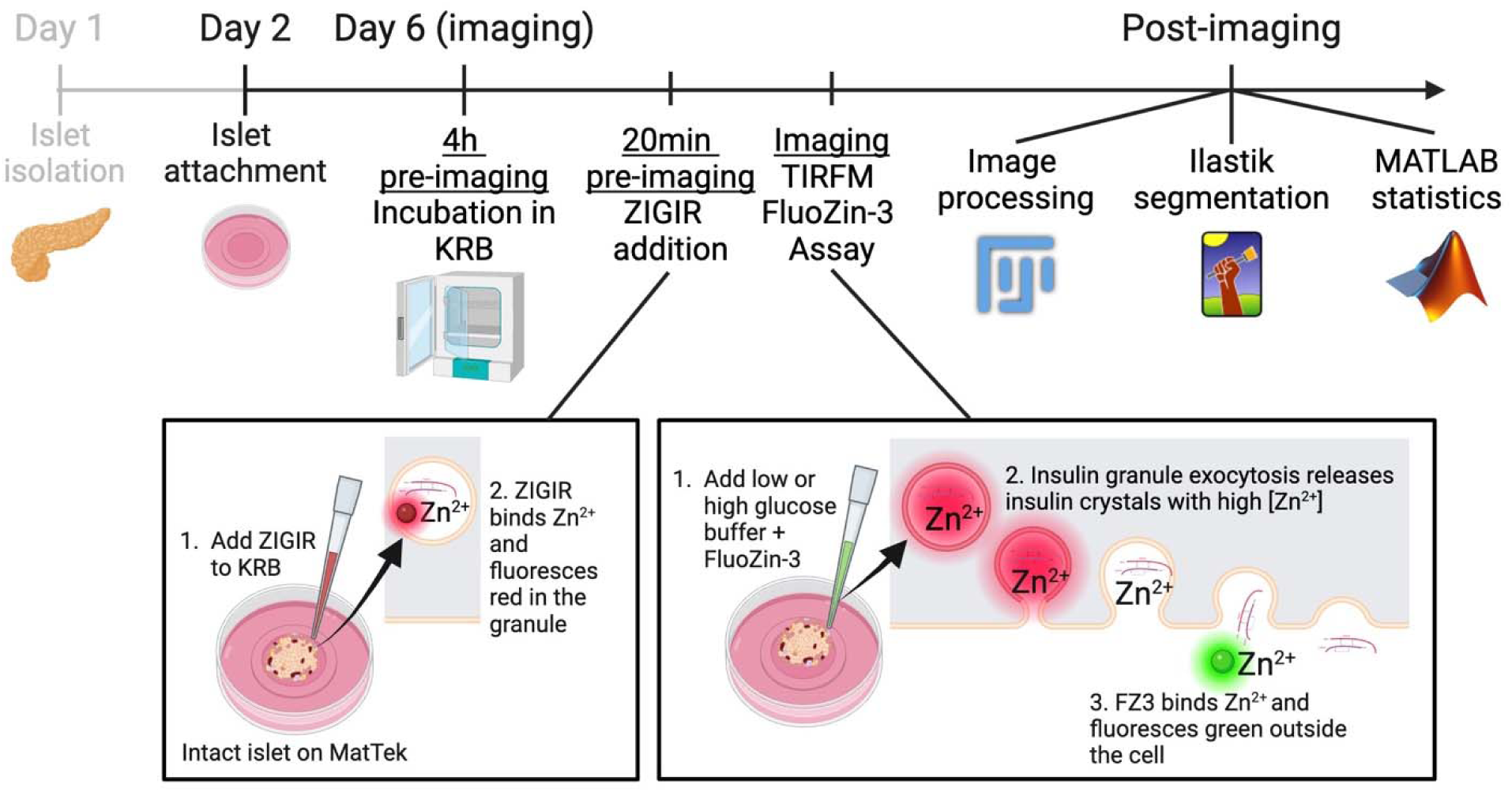
Schematic of the workflow from islet isolation to image processing and analysis. Islet isolation and attachment is detailed in Ho et al. 2023^18^. Islets are attached three to five days prior to imaging. On day of imaging, islets are incubated in low glucose (2.8mM) KRB for two to four hours. ZIGIR is added 20 minutes prior to imaging. FluoZin-3 is added at the time of imaging. Post-imaging processing and analyses include 1) a pre-Ilastik workflow to heighten signal-to-noise of flashes; 2) the Ilastik workflow to automatically detect flashes; and 3) the optional MATLAB script to detect clustering of granules.

### Islet isolation and plating

Refer to Ho et al. 2023^18^ for a detailed protocol for islet isolation. Following islet isolation and appropriate period of recovery for freshly isolated islets, attach islets to appropriate ECM three to five days prior to imaging. This step is critical for allowing full attachment and preventing premature islet detachment during imaging, especially during addition of any treatment solutions.

### Islet Imaging

After incubating islets for at least 2 hours in low glucose KRB (2.8mM), cells will be ready to image. If using the ZIGIR addition, incubate in ZIGIR for 20 minutes prior to imaging. Also use any pre-incubations at this time such as drugs of interest. Using TIRF microscopy, focus islets in the TIRF plane with special attention to the fluorophore of choice. The use of TIRF microscopy for this protocol is critical as events are difficult to see without the high signal-to-noise ratio that TIRF allows for. For example, traditional laser scanning and spinning disk confocal systems are not sufficient for imaging this phenomenon. Add FluoZin-3 dye concurrently with glucose to stimulate secretion. Treatments of interest may also be added at this time for shorter-term assessment.

### Processing and Analysis

Movies will need processing to divide the image by itself and obtain a low signal-to-noise movie emphasizing the flashes. These movies can then be loaded into the Ilastik programs for pixel classification and subsequently object classification. The pixel classification yields prediction maps for the flashes, while the object classification program yields binary decisions from the machine learning based on those probability maps. Therefore, both are required for complete flash detection in this analysis. Resulting coordinate data from Ilastik, alongside cell borders, can next be loaded into the custom MATLAB script to further assess insulin secretion clustering.

### Limitations

This technique is optimal for assessing solely first phase insulin secretion for two reasons. 1) The ability to capture rapid events necessitates continuous imaging at a frame rate of at least 10 frames/second. Continuing to image and expose islets to laser for longer than this 10 minutes may result in unwanted phototoxicity and/or photobleaching. 2) Although these flashes are transient, the increasing extracellular [Zn^2+^] increases background signal as the movie goes on. Again, this increasing background signal is sufficiently low to allow for clear imaging for the duration of first phase secretion, but hours of imaging to recapitulate second phase secretion may make detection of these events difficult. For these reasons, it is recommended to image first phase secretion only.

This protocol also does not allow for full specificity of secretion events. Although we can be reasonably confident in observing only insulin secretion events on account of glucose stimulation, alpha cells also have glucagon granules containing Zn^2+^. For this reason, we cannot rule out that some flashes may be alpha cells secreting glucagon. However, the likelihood of this upon glucose stimulation is minimal and if using a cell line with a beta cell marker like our H2B-Ins-mApple marker, cells that are not beta cells can be ruled out of this analysis.

Finally, the combined ZIGIR/FluoZin-3 technique presents an interesting limitation in that ZIGIR does not appear to label all granules. This can be observed in our data during which some secretion events happen but there is no clear insulin granule disappearance from inside the cell. This may be due to lack of available binding for ZIGIR to zinc in tightly packed granules. However, between using both the FluoZin-3 and the ZIGIR assay, we are likely to account for all granules we are interested (that is, those that secrete).

## Materials

### Biological Materials

Islets or cells of choice.

CRITICAL: We find that this assay works best for intact mouse islets, but it has also been performed using the MIN6 beta cell line as well as dissociated islets, which have been lightly digested to break them down into cell clumps.

### Reagents

1. Invitrogen™ FluoZin™-3, Tetrapotassium Salt, cell impermeant (Invitrogen, F24194), reconstituted to 5mM in MilliQ H_2_O CRITICAL: this reagent must be the cell impermeant tetrapotassium salt, as the cell permeant version of FluoZin-3 will act similarly to the ZIGIR and be unable to label extracellular secretions.
2. ZIGIR (VitalQuan, 0143), reconstituted to 1mM stock and used at 0.1µM working concentration.
3. CellMask Invitrogen, C10046) used at 2.5 µg/ml from 5mg/ml stock concentration.
4. Krebs-Ringer Bicarbonate (KRB) Buffer (see below for reagent setup for low glucose and high glucose KRB compositions)

CRITICAL: KRB is necessary in this assay for multiple reasons. It is an important standard for glucose stimulated insulin secretion assays, keeping islets at physiological concentrations of ions to ensure survival during assays; allowing for precise control over glucose level; and importantly, in this assay, removing any extracellular Zn^2+^.

### Equipment

1. Nikon TE2000E Microscope with Nikon TIRF2 System for TE2000
2. TIRF 100X 1.49 NA oil immersion lens
3. Andor iXon EMCCD camera
4. Nikon Elements C software
5. TOKAI HIT WSKM Sample Temperature Feedback Heating Stage Top Incubator INUBTFP-WSKM with dish attachment for 35mm dish
6. Lens cleaner and lens paper
7. Micropipettes capable of 0.2µl and 38.8µl volumes
8. Fiji^28^
9. ilastik^17^ (Version 1.4.0.post1)
10. MATLAB (Version R2024a, The MathWorks, Inc.)

### Reagent Setup

FluoZin-3: Best if mixed fresh, immediately prior to imaging, on ice. Dilute to 20µM in desired glucose concentration of KRB (low or high glucose). It is best to do this under a lamp near the microscope, so you can transfer the dye quickly but also so you can see that you have actually pipetted what may need to be a small volume of FluoZin-3. Mix by pipetting, then load the pipette tip with the solution and rest it nearby so you can quickly retrieve it. Add to islet dish when ready.

ZIGIR: Best if pre-incubated for 20min-1 hour at room temperature, depending on imaging system and desired fluorescence intensity. From a stock of 1mM, dilute to 0.1µM directly in the islet dish. Wash briefly in same KRB composition prior to imaging to reduce background fluorescence by gently aspirating and replacing the volume.

CellMask: Prepare in KRB buffer at 2.5 µg/ml from a 5mg/ml stock. Post-FluoZin-3 imaging, add 50µl of the 2.5µg/ml to the dish and allow to incubate for 10-15 minutes. Image at 642 nm by adjusting the TIRF angle as needed.

Low glucose (2.8 mM) KRB: recipes for stock solutions and recipes for various volumes of KRB. Add in the following order:

| Reagent/Desired Volume | 50mL (in µl) | 30mL (in µl) | 20mL (in µl) | MW (g/mol) | Stock |
| --- | --- | --- | --- | --- | --- |
| MilliQ water | 40.5ml | 24.3ml | 16.2ml | n/a | n/a |
| 2.5M CaCl<sub>2</sub> | 46 | 27.6 | 18.4 | 110.99 | 2.744g in 10ml H<sub>2</sub>O (difficult to dissolve) |
| 1M NaCl | 5550 | 3330 | 2220 | 58.44 | 14.61g in 250ml H<sub>2</sub>O |
| 1M KCl | 240 | 144 | 96 | 74.56 | 3.728g in 50ml H<sub>2</sub>O |
| 1M MgSO<sub>4</sub> | 60 | 36 | 24 | 120.37 | 6.02g in 50ml H<sub>2</sub>O |
| 1M KH<sub>2</sub>PO<sub>4</sub> | 60 | 36 | 24 | 136.09 | 6.81g in 50ml H<sub>2</sub>O |
| 0.5M glucose | 280 | 168 | 112 | 180.16 | 4.5g in 50ml H<sub>2</sub>O (filter sterilize) |
| 0.5M HEPES | 1000 | 600 | 400 | 238.30 | 5.96g in 50ml H<sub>2</sub>O (filter sterilize) |
| 10% BSA | 1000 | 600 | 400 | n/a | 5g in 50ml H<sub>2</sub>O (store in aliquots at -20°C) |
| 1M NaHCO<sub>3</sub> | 1250 | 750 | 500 | 84.01 | 4.2g in 50ml H<sub>2</sub>O (takes 2-4h to dissolve at RT) |

For high glucose (25mM) KRB, it is recommended to create a stock of 50mM by adding 1:10 0.5M glucose:low glucose KRB (e.g. 900µl low glucose KRB + 100µl 0.5M glucose). Then, add volume of 50mM glucose KRB to the 2.8mM glucose KRB to get your final desired concentration of glucose, whether that is 11, 16.5, or 25mM (e.g. if islets are in 50µl media, add 50µl of 50mM glucose KRB to get a final concentration of 25mM KRB).

### Procedure

#### Step 1, Islet Attachment

##### 3-5 days prior to imaging (20 minutes)

1. Three to five days prior to imaging, plate cells onto a plasma-cleaned vascular extracellular matrix-coated one-chamber 10mm MatTek dish. See Ho et al. 2023^18^ for clear instructions on this procedure. Use whatever transfection, transduction, cell density, etc. you are looking to analyze.

a. CRITICAL STEP: the longer that islets are left attached to the dish, the flatter they will become. However, the shorter they are left attached, the higher the probability they will detach during imaging and manipulation. For this reason, the recommended optimal time of attachment is 4 days.
b. TROUBLESHOOTING: If islets are not attaching well or detach easily, it is recommended to increase attachment time. However, this may require addition of extra media and will increase the flattening of the islet, which could disrupt some cellular processes.

#### Step 2, Imaging

##### 2.4. hours of pre-incubation in KRB, ∼10 minutes per dish for imaging

1. Make low glucose (LG; 2.8mM) and high glucose (HG; 50mM) KRB fresh on the day of imaging. See recipe below.
2. Gently aspirate the media from the dish using a pipette, not the vacuum line, and add 100µl of LG KRB.

a. CRITICAL STEP: using the vacuum line may detach islets. Avoid excessive aspiration to prevent islet detachment.
b. Halfway through incubation period, gently aspirate and replace with another 100µl of KRB. The final volume prior to imaging should be 100µl.
c. Following this replacement of KRB, ZIGIR can be added if desired. Add 0.1 µl of stock of 1mM to 100 µl of KRB and from this, add 10 µl to dish to get a final concentration of 0.1µM, approximately 20min-1hr before imaging.
d. TROUBLESHOOTING: Longer incubation times will increase fluorescence, but longer than 20 minutes is not required for thorough IG labeling. The concentration of ZIGIR can also be adjusted and optimized for individual systems.
3. Prior to beginning imaging, retrieve an aliquot of stock 5mM FluoZin-3 from the freezer and keep on ice for the duration of imaging.
4. Before mounting the dish on the microscope, gently clean the bottom of it with lens cleaner and lens paper.

a. CRITICAL STEP: This is to reduce random particles that disturb the optical path. Be sure to avoid detaching the islets during this process.
5. Find islet(s) using brightfield and save the position(s). Perform any further microscope optimization necessary for your microscope at this time.
6. Acquire a brightfield stack ∼10µm into the islet and any further images you require at this time (Figure 2a-c).

a. CRITICAL STEP: The brightfield stack is critical for assisting in outlining cells during analysis. This step can also be done following the experiment if desired, but some fluorescence may need to be acquired prior to FluoZin-3 (e.g. other green fluorophores).
b. Optionally, use CellMask dye (2.5 µg/ml, Invitrogen, catalog. C10046) to label the islet membrane following movie acquisition, to more faithfully outline the cell boundaries.
7. Set the critical angle (and direction if not ring-TIRF). This will likely need to be performed every time due to slight differences in MatTek dishes and samples.

a. TROUBLESHOOTING: An ideal method for setting the critical angle is to test several (on low laser power to avoid photobleaching/toxicity) and choose the lowest angle that still allows for signal and high signal-to-noise ratio. Individual systems may vary.
8. Prepare acquisition settings as appropriate for your experiment, and set up a time lapse with no delay for 6-10 minutes. An exposure time of 30-50ms is preferred to obtain a final frame rate of 10-15 frames/second.

a. TROUBLESHOOTING: We find that at these settings, a laser power of 10-30% on our system is adequate. Less if there are no other fluorophores in the sample, more if there are other fluorophores present. Individual systems may vary.
9. Once the sample is mounted and acquisition settings are set, prepare the FluoZin-3 solution by adding 38.8µl of LG or HG KRB (depending on your desired stimulation). Add 0.2µl of 5mM stock FluoZin-3, for a final working concentration of 20µM.

a. CRITICAL STEP: It is best to do this under a lamp near the microscope, so you can transfer the dye quickly but also so you can confirm that you have actually pipetted the FluoZin-3, as it can be difficult to see.
10. Mix by pipetting, then load the pipette tip with the 40µl FluoZin-3 solution and rest it nearby so you can quickly retrieve it.
11. Begin acquisition and carefully remove the incubating chamber lid, being extremely careful not to disturb the movie. Add FluoZin-3 solution dropwise to the center of the dish, without creating bubbles (Figure 2d-f, Supplemental Video 1).

a. TROUBLESHOOTING: the entirety of the field will begin to turn green due to increasing extracellular [Zn^2+^], but if pixel intensity becomes saturated and signal is “blown out,” you can consider reducing laser power or the FluoZin-3 final concentration for optimization.
12. Real-time secretion is visible as a circular flash that lasts ∼100ms or 1-2 frames (depending on exposure time and speed of camera; Figure 2g-j, Supplemental Video 2). IGs with ZIGIR are visible as circular granules 200-300nm in size that have longer-lasting dynamics (Supplemental Video 3).

a. Critical: these are difficult to detect by eye in raw data, but will become clearer after processing.
13. Following acquisition, CellMask may be added (50µl from 2.5 µg/ml of working concentration) and acquired to better outline cells during processing.

#### Processing and Analysis

1. Open the movie in ImageJ/Fiji. If necessary, separate channels so that you are only working with the green channel in which you have acquired FluoZin-3.
2. Duplicate the stack without the first slice (e.g. if it is 1-5000 frames, duplicate as 2-5000). This is stack A.
3. Again duplicate the original green channel, this time without the last slice (e.g. duplicate 1-5000 as 1-4999). This is stack B.
4. Create a divided movie by using the image calculator in Fiji under the Process menu and divide stack A/stack B. Make sure “create 32 bit” is checked. Save this as the “processed” movie.

a. The resulting movie will look like static, but any flashes will be circular white regions. It is recommended to set the lookup table to grays to enhance contrast (Figure 2d2-f2, Figure 3a).
5. We have developed a Fiji macro entitled “Pre-ilastik processing” to complete steps 2-4 (see Supplementary Software)
6. Following the creation of the processed movie, there are 2 options for identifying flashes. The first is manually, by using the single point tool and ROI manager to add each point individually. This process is time-consuming but allows you more control over what you deem to be a flash or not. This process is as follows:

a. Use the point tool to manually identify flashes (set your own personal criteria for brightness, shape, etc.).
b. Add each point individually to the ROI manager.
c. Under the Analyze menu, choose Set Measurements so that you only get the Xy coordinates and stack position.
d. Save the file of ROIs, then export these points using “measure” to measure all from the ROI manager. You should only have 4 columns: ID, X coordinate, Y coordinate, and stack position.

i. The number of columns is important for inputting into the MATLAB script for cluster analysis.
e. Convert the .csv to an Excel Workbook. This step is also critical for the MATLAB script to read the data.
f. Save the file as “Flashes”. CRITICAL: This nomenclature is also critical for the MATLAB script to read the data.
7. The second option is to use the custom machine learning program developed in our lab created using the Python-based machine learning freeware ilastik. The following instructions pertain to the automated flash identification using ilastik:

a. Export your processed movie as an .hdf5 (.h5) using the Plugins menu>ilastik>Export HDF5. This compresses the movie in a format appropriate for ilastik. ilastik will take .tif/.tiff, but works much better on .hdf5 (Figure 3a).
b. Install and open ilastik.
c. Open the ilastik pixel classification program linked, entitled “FluoZin-3 Pixel Classification Step 1”. This program comes ready to compute experimental data and there is no need to add any further training data to your “input data.” TROUBLESHOOTING: However, if you find that the program does not accurately identify your FluoZin-3 flashes, possibly due to slight differences in acquisition, you may want to add additional training files to “input data” (Figure 3b-b1).
d. If you choose to add more training data, it is recommended to use 5-10 frames of your movie that have flashes as a positive control example for ilastik. You can use full-length movies, but this will be memory-intensive for ilastik and your computer. Steps for training can be found here: https://www.ilastik.org/documentation/pixelclassification/pixelclassification. Make sure these training movies are also in .hdf5 format.
e. CRITICAL: when you start training, if you use “store absolute path” for the file location, you MUST keep those files you input EXACTLY where they are. Moving them to even a sub-folder of the folder they are in can disrupt the path and render the ilastik program unopenable. Instead, double click on the “location” of the file and change it to “copy into project file.” This will allow you to move those files around without penalty.
f. Once you feel your training is sufficient or you do not need more training, batch process your full-length processed movies. Do not exit ilastik during this time. Not only will you lose data progress butt you may corrupt the program file. It is also strongly recommended to not force quit ilastik any time for the same reason.
g. Export any training and/or experimental data as .hdf5 files. This should result in files with a “pixel classification” suffix (Figure 3b2).
h. Now open the object classification ilastik program linked, entitled “FluoZin-3 Object Classification Step 2”.
i. Input any of your raw .hdf5 training data (Figure 3a) and their corresponding pixel classification .hdf5 files (Figure 3b2) if you need to train the object classifier further, similarly to the pixel classification workflow (Figure 3c-c3).
j. Add your raw .hdf5 full-length processed movies and their corresponding pixel classification .hdf5 files to batch processing.
k. You may wish to export some of this data as an Excel file; to do so, set image export to include the .csv. This file is helpful because it contains coordinates of the center of shapes and can help you skip a step in Fiji (Figure 3c3).
l. To open the object classification files in Fiji, use Plugins>ilastik>Import HDF5 (it won’t work to drag and drop with bio-formats or open from the file menu).
m. To check and ensure that the flashes layer has been imported, change the LUT using Image>Lookup Tables>glasbey (this is the only LUT that will allow you to see the binary classifications; Figure 3d).
n. Save this as a .tif/.tiff for easy access.
o. . Threshold the image so that the flashes are included in signal (Figure 3d1):

i. TROUBLESHOOTING: ilastik may modify the output such that if you threshold normally, it turns the flashes into pixel value=0 and the background into pixel value>0. To fix this, go to Edit>Options>Appearance and check “use inverting lookup table” to invert the background and flash values.
p. Proceed to thresholding with the following selected:

i. Default, over/under > check “dark background” and “don’t reset range” > apply
ii. Under “apply” pop-up menu, set Method: default, Background: dark, and check “black background” and “create new stack”.
q. Now use Analyze>Analyze Particles to identify flashes and their (x,y) coordinates. Again, ensure that the measurements are set appropriately. Use the following parameters:

i. Size: 0-Infinity
ii. Circularity: 0-1
iii. Show: Outlines
iv. Display results
v. Add to manager
vi. Exclude on edges
r. To obtain center coordinates, Set measurements to include “centroid” and “stack position.”
s. In Fiji with all of the flashes open, click “measure” in the ROI manager to get the (x,y) coordinates of the flashes. These can be plotted with the macro “multi point plotter” (Figure 3d2-3).
t. CRITICAL: Depending on the signal-to-noise ratio of your movie, you may need to manually delete some flashes that land outside of the islet.
u. CRITICAL: Depending on the frame rate of your movie, you may also need to manually confirm that there are no instances of two flash outlines per flash (e.g. at frame 1 and 2, 100ms apart).
v. Some flashes will be smaller or larger and will have higher or lower accuracy, respectively (Figure 3e-f), but show a much closer estimation of the midpoint than manual annotation (Figure 3e3-f3).
w. Save these as an ROI and proceed depending on your analysis of interest.
x. We have developed a Fiji macro entitled “Post-ilastik processing” to complete steps m-v (see Supplementary Software).
8. At this point, if you are interested in identifying clusters with the MATLAB program (Supplementary Software “MATLAB Secretion Clustering Program”), use the following steps to properly export necessary data:

a. Set measurements to include “centroid” and “stack position.”
b. In Fiji with all of the flashes open, click “measure” in the ROI manager to get the (x,y) coordinates of the flashes.
c. This should give a text file with only four columns: the ID (unlabeled column), X coordinate, Y coordinate, and Slice.
d. CRITICAL: Convert the .csv to an Excel Workbook. This step is also critical for the MATLAB script to read the data.
e. CRITICAL: Save the file as “Flashes”. This nomenclature is also critical for the MATLAB script to read the data.
f. Next, define cells for MATLAB. Outline the islet, cells, or regions as appropriate, depending on your analysis of interest, using brightfield (Figure 4a) or CellMask (figure 4b). With a single of these ROIs selected, use Analyze>Tools>Save XY coordinates. Do this for each ROI as needed (Figure 4c).
g. The resulting Excel .csv should only have 3 columns: X, Y, and Value. (Note, the ROI measurements MUST be in a .csv, contrary to d) (Figure 4d).
h. CRITICAL: Save this in the format “ROI#” where # = whatever number value you assign to that particular ROI. This nomenclature is also critical for the MATLAB script to read the data. (E.g. if you have 50 cells, you will want ROI1 through ROI50 in a folder, each as separate files).
i. CRITICAL: Make sure all these files are in the same directory together and run the MATLAB script (see Supplementary Software, entitled “MATLAB Secretion Clustering Program”). These two components, flashes (Figure 5a) and cell outlines (Figure 5b) must be properly annotated for MATLAB input.
j. The MATLAB script should yield an ROI map with color-coded clusters (Figure 5c) and an Excel file with information on these clusters (Figure 5d-f, for more see Trogden et al. 2021^10^).

i. TROUBLESHOOTING: if coordinates of flashes or ROIs appear flipped, use Analyze>Set Measurements and check “Invert Y coordinates” and re-export everything and re-run the analysis. This may be due to an orientation setting in the microscope acquisition software or in Fiji.
ii. From here, statistics regarding per-cell secretion in clusters or not can be used for further analysis (Figure 5g-h).

### Troubleshooting

Islet attachment is one of the most critical steps in this protocol and may also be one of the most difficult. Detachment can occur at any point during the process, particularly if attachment is insufficient, and renders the islets unusable for this assay. To avoid detachment, optimization of the islet attachment protocol is critical. This may include altering dish plasma cleaning parameters, amount and distribution of ECM used, time to allow ECM to “cure” before adding, duration of attachment, addition of media, etc. It must also be noted that the ECM used here has been discontinued and replaced with a Matrigel mixture of similar composition. This may require its own optimization.

**Figure 2.**
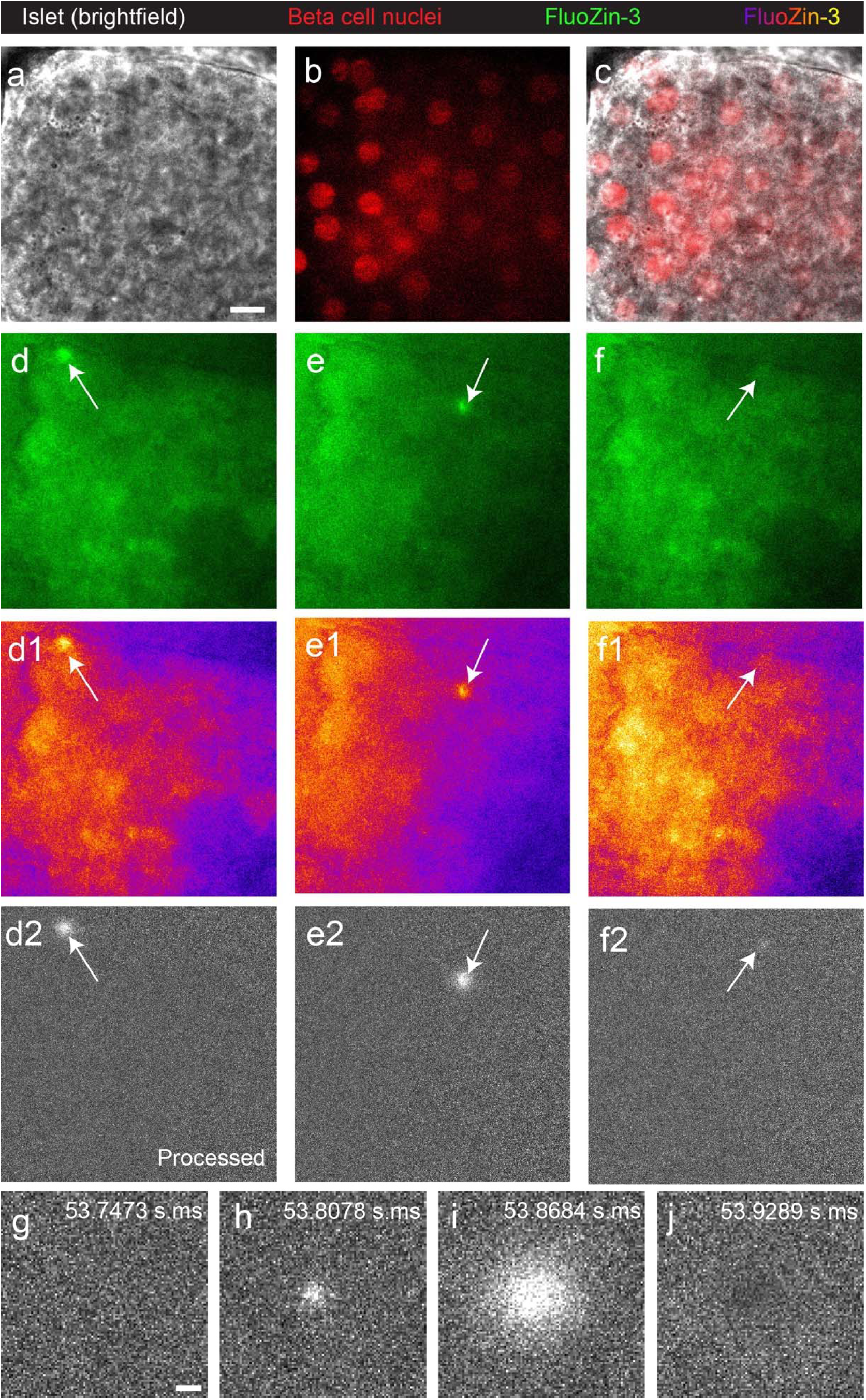
The FluoZin-3 assay resulting images and processed images. a) Intact mouse islet imaged on TIRF using brightfield illumination. b) Beta cell nuclei identified using the beta cell-specific H2B-Ins-mApple probe. c) Membrane outlines using a combination of the membrane probe, brightfield image, and beta cell nuclei. d) Example flash of FluoZin-3 labeling an exocytosis event, white arrow, in the raw file, d1) in the raw data with a fire LUT to show intensities, and d2) in the processed data with divided background. e-e2) Another example of a flash, white arrow, in the darker side of the illumination plane. f-f2) An example of a flash, white arrow, with a low signal-to-noise ratio, that is difficult to see in f-f1) the raw data, but becomes visible upon f2) processing. Scale bar for all = 10 µm. Corresponds to Supplemental Video 1. g-j) Montage of a single flash over time at g) 53.7473, h) 53.8078, i) 53.8684, and j) 53.9289 m.ms, illustrating the transient nature of the flashes. Scale bar = 2 µm. Montage corresponds to Supplemental Video 2.

**Figure 3.**
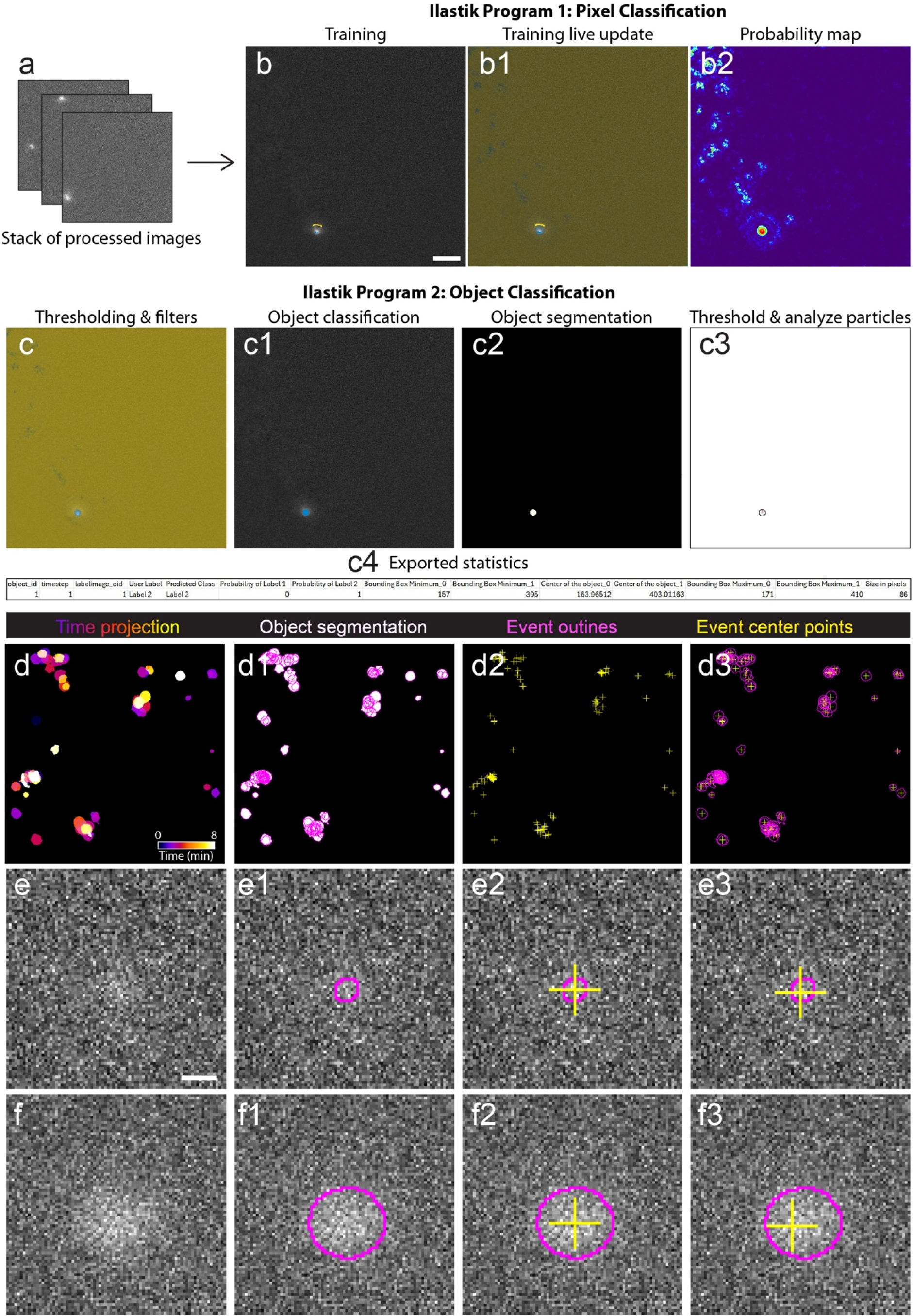
The Ilastik workflow input, processing, resulting, and processed images. a) The input for the first Ilastik program the Pixel Classification workflow, is a stack of images that have been background division processed as in Figure 2d2-f2, that has been converted to a .h5/.HDF5 file. b-b2) Representative images of a single slice in the Ilastik Pixel Classification workflow. b) Training involves small annotations (yellow and blue drawing) to identify two labels, followed by b1) live updates to confirm annotations are updating the prediction maps. b2) A probability map that has been exported for this slice and visualized in Fiji using the thermal LUT to demonstrate the pixel prediction map which will get fed into c-c4) Ilastik program 2, the Object Classification workflow. c) Thresholding and filters visualizes the parameters that will go into final object classification, using the pixel prediction maps. c1) Object classification consists of manual identification of objects from each label. c2) An exported object classification file that has been imported into Fiji shows the object segmentation and can here be thresholded and made binary. c3) After thresholding and binarization, Analyze Particles can be used to identify flashes. c4) Displays the statistics from this single flash, importantly including the center of object statistics. d-d3) Post-Ilastik processing. d) Time projection using the fire LUT in Fiji, showing the Ilastik-identified flashes over time (note that this is not a required step, but included for illustrative purposes). d1) Object segmentation as in c3, white, including the outlines of identified flashes (magenta). d2) Center points of the identified the objects (yellow crosses). d3) Outlines (magenta) and center coordinates (yellow crosses). Scale bar for all = 10 µm. e-e3) Example of a relatively smaller flash with high-confidence center coordinate detection. e) Example small flash. e1) Outline of segmented flash by Ilastik (magenta). e2) More precise, automatic center coordinate detection by Ilastik (yellow cross). e3) Less precise, manual center coordinate detection (yellow cross). f-f3) Example of a relatively smaller flash with lower-confidence center coordinate detection. f) Example large flash. f1) Ilastik outline (magenta). f2) More precise, automatic center coordinate detection by Ilastik (yellow cross). f3) Less precise, manual center coordinate detection. Scale bar for all = 2µm.

**Figure 4.**
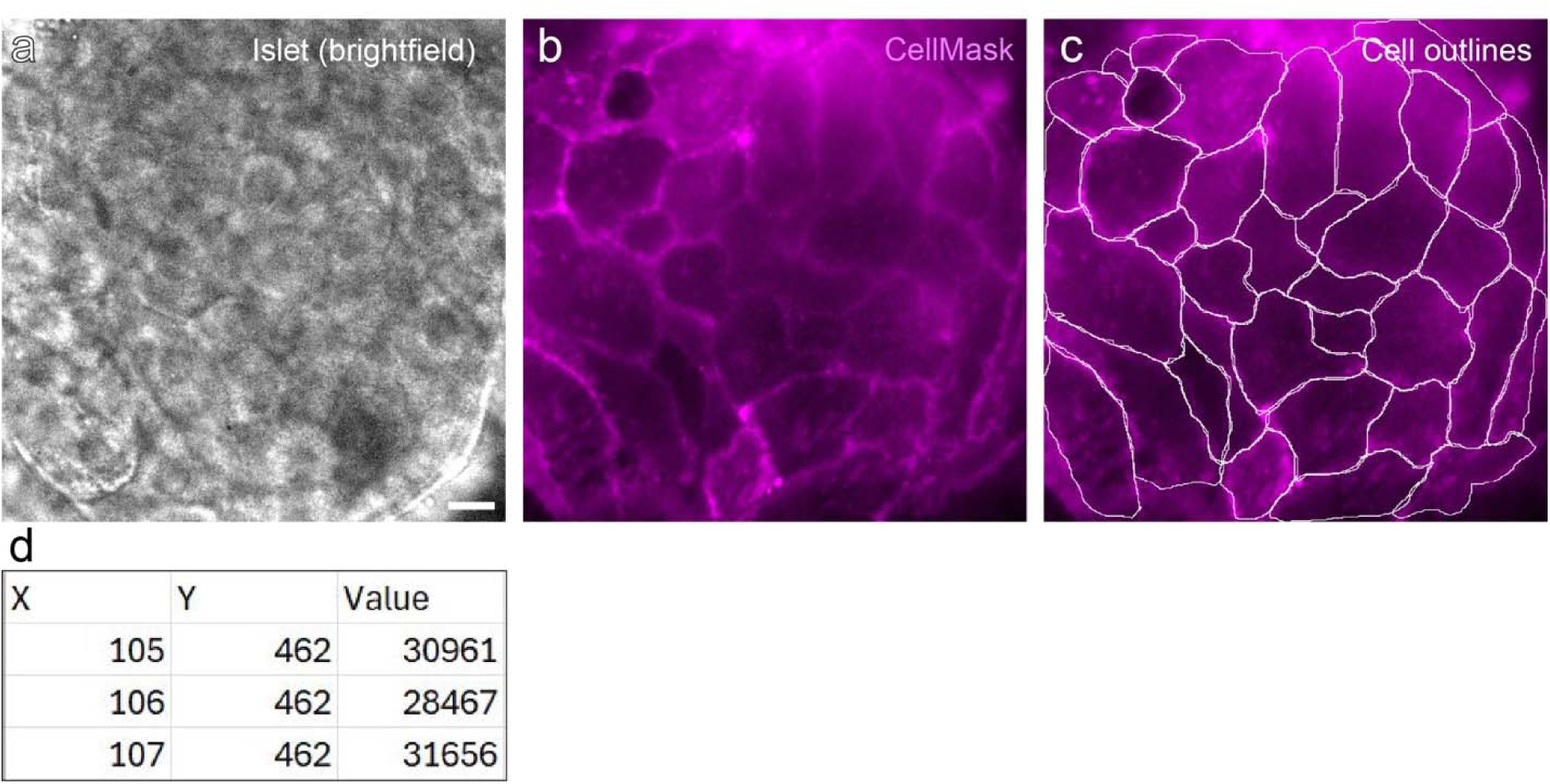
Cell definition guided by brightfield or CellMask. a) Brightfield image of an intact islet, taken prior to imaging, acquired as a stack ∼10µm into the islet. b) CellMask added to the same intact islet post-FluoZin-3 assay, acquired as a single slice at the bottom of the islet. c) Cell outlines drawn according to the CellMask image. Scale bar for all = 10µm. d) Example of the spreadsheet columns required in a single cell ROI for input into the MATLAB script.

**Figure 5.**
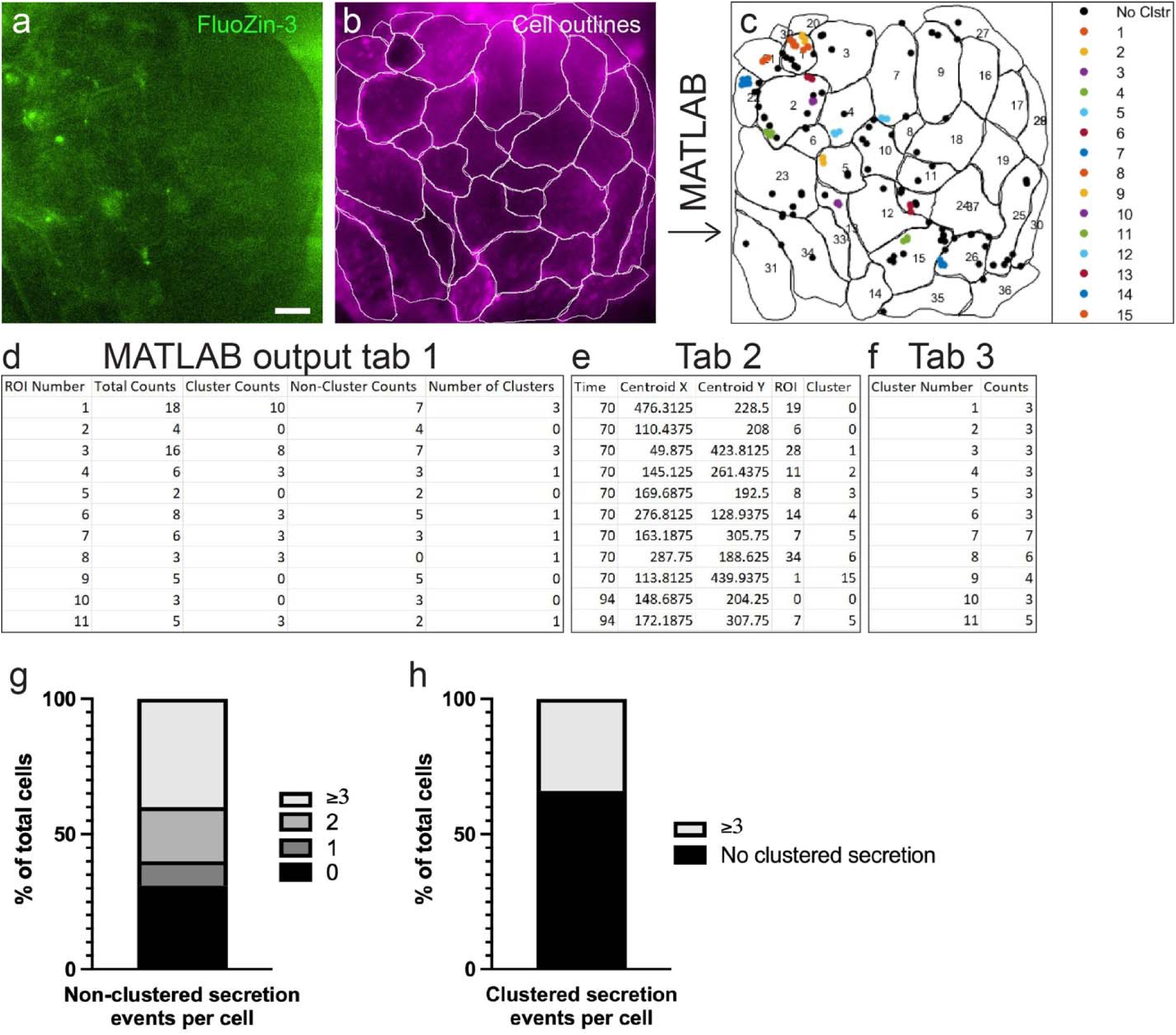
MATLAB clustering analysis workflow and example of statistics from a single islet. a) FluoZin-3 MaxIP of a few frames, exemplifying flashes. b) Cell outlines using CellMask, as in Figure 4b-c. Coordinates from a) and b) constitute the input for the MATLAB program which outputs c-f). c) MATLAB-generated figure showing cell outlines and their IDs, and secretion events coded by cluster number (non-clustered events = black, clustered events = color; right). Scale bar for all = 10µm. d) MATLAB-generated spreadsheet tab showing per-cell statistics: the total secretion events (“Total Counts”) per cell (“ROI Number”, corresponding to cell identities in c)), secretion events part of a cluster (“Cluster Counts”), secretion events not part of a cluster (“Non-Cluster Counts”), and Number of Clusters. e) Tab showing time-based statistics: frame at which the event occurs (“Time”), xy coordinates of the secretion event (“Centroid X,” “Centroid Y”), cell the event belongs to (“ROI”), and cluster ID the event belongs to (“Cluster”). f) Tab showing per-cluster statistics: cluster ID (“Cluster Number”) and number of events within that cluster (“Counts”). g) Plotted statistics for the single exemplified islet depicting percent of total cells with no non-clustered secretion events (“0”, black), percent of cells with one non-clustered secretion event (“1”, dark gray), percent of cells with two non-clustered secretion events (“2”, medium gray), and percent of cells with 3 or more secretion events (“_≥_3”, light gray). h) Similar descriptive statistics, but for clustered secretion. Depicting percent of total cells with no clustered secretion events (black) or 3 or more clustered secretion events (“_≥_3”, light gray).

Fluorescence optimization may also be necessary for this protocol. As mentioned above, the ZIGIR dye is quite bright and stable, and may need to be reduced in concentration and pre-incubation time to prevent over-exposure and bleed-through to the green channel. As described previously^14, 15^, 15-20 minutes is acceptable for labeling the majority of granules, and longer incubation times likely do not increase labeling. However, optimization for increased fluorescence may include longer incubation times and higher concentration. This may differ significantly (on the scale of nM-µM) for cell lines versus intact islets versus tissue. Additionally, ZIGIR penetrates the outer layer of the islet quite well, but it becomes more difficult in subsequent layers for penetration to occur, despite its ability to traverse membranes. Optimization here may be needed. Setting the optimal TIRF critical angle and laser power(s), dependent on the system being used, may also help to optimize this concentration. Additionally, it is noted above that as insulin secretion progresses throughout first phase, the entirety of the field may fluoresce due to increasing extracellular Zn2+ concentrations. This may result in pixel saturation of the field, in which case a combination of optimization between FluoZin-3 concentration, laser power, and TIRF angle may be appropriate for preventing this issue.

The existing Ilastik programs have been custom-trained for images acquired on our system. There is a great deal of variation between systems and even between identical microscope rigs in different locations. For this reason, it may be necessary to add or replace training files in the Ilastik workflow and custom train for the most optimal segmentation of FluoZin-3 flashes. Several other differences between systems may mandate further optimization and customization of the processing and analysis workflow described herein.

### Timing

As described above, the main constraints on time in this protocol are between islet attachment and desired time of imaging. It is recommended to decide on a desired time of imaging first, which requires the attachment to take place 3-5 days beforehand, and islet isolation from mice to occur between one day to two weeks prior to attachment. Processing and analysis can be performed at any time subsequent to imaging.

### Anticipated Results

This protocol should generate live insulin secretion movies with labeled insulin granules with precise localizations of secretion sites and correlation between secreted granules and disappearing live insulin granules. Specifically, the movies generated should be single or dual color in green or red and green channels. The frame rate should be no less than ten frames per second to adequately capture secretion events in their entirety. We include here a full-length movie of flashes that has worked well, along with a movie that does not secrete well and a low glucose stimulated movie which should display no flashes. We also include images of ZIGIR dye that display excellent islet penetration as compared to sub-optimal penetration.

The processing workflow of these movies will also yield detailed statistics regarding the flashes as a result of the ilastik processing, object segmentation that can be visualized in Fiji, and clustering analyses with detailed statistics. We also show in Figures 3 and 5 examples of these processing spreadsheets, object segmentation files, and clustering analyses.

These results can be used to perform shortest distance measurements as in Fye et al. (2025)^16^, co-localization analyses, etc. as desired.

Using this methodology we analyzed the insulin secretion events of different IG pools labeled by ZIGIR (Figure 6a): predocked, which appear before high glucose stimulation and are docked at the membrane (Figure 6b); docked, which appear upon high glucose stimulation and dock at the membrane; and newcomer, which appear upon high glucose stimulation but don’t dock at the membrane (Figure 6c). We obtained the precise subcellular location of these insulin secretion events in beta cells using cell boundaries determined using CellMask dye or brightfield images (Figure 4). We analyzed the behavior of secretion events (non-clustered vs clustered secretion events; Figure 5), and found that 31% of cells do not secrete while 69% of cells displayed 1, 2 or ≥3 non-clustered secretion events (Figure 5g). On the other hand, we found only 33% of cells secrete in clusters of ≥3 secretion events (Figure 5h), indicating the secretory heterogeneity of islet beta cells. This method is useful in the investigation of functional beta cell heterogeneity of IG secretion in space and time.

**Figure 6.**
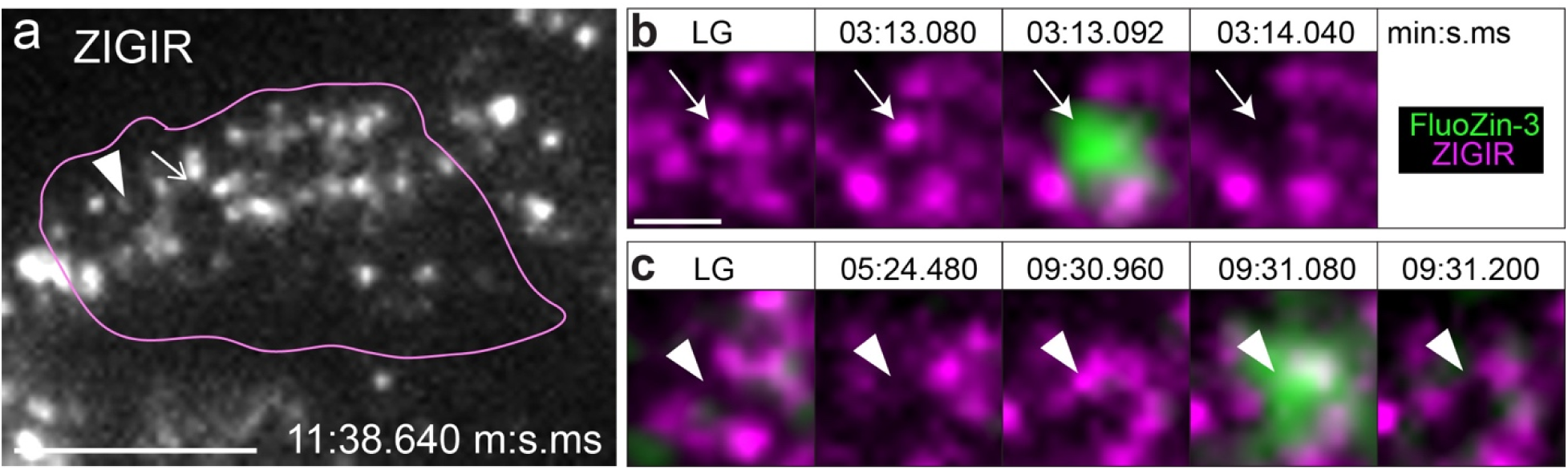
FluoZin-3 and ZIGIR reveal behavior and dynamics of secreted IGs. a) ZIGIR (grays)-labeled granules in a single cell (magenta, outline). Right arrow corresponds to predocked IG, b). Left arrowhead corresponds to newcomer granule, c). Scale bar = 5µm. b) Example of a predocked granule at the membrane in low glucose (LG), which gets secreted at 03:13.092 min:s.ms. c) Example of a newcomer granule which is not at the membrane in LG nor earlier in the movie at 05:24.480 min:s.ms but arrives at 09:30.960 min:s.ms and gets secreted thereafter at 09:31.080 min:s.ms.

## Supporting information

Pixel and Object Classification Files

FIJI MACROS

MATLAB Secretion clustering program

## Abbreviations

(IGs): Insulin granules
(TIRF): total internal reflection fluorescence
(LG): low glucose
(HG): high glucose
(KRB): Krebs-Ringer Bicarbonate

## Acknowledgements

This protocol was adapted from Zhu et al. (2015) and Trogden et al., (2021) and is a continuation of the protocol laid out in Ho et al. (2023). This work was supported by National Institutes of Health (NIH) grants F31 DK13344-01 and T32 DK007563-33 (to MAF; O’Brien, PI), NIDDK R01 DK106228 MPI (to IK and GG), NIGMS MIRA R35GM127098 (to IK), R01-DK65949 (to GG), and R01-DK125696 (to GG). The authors would like to thank Hamida Ahmed for crucial technical support and the Kaverina lab for critical review of the manuscript.

## Author contributions

I.K. was responsible for the intellectual development of the procedure. G.G. facilitated mouse work and contributed to intellectual development of the procedure. M.A.F. and R.S. equally contributed to generating figures and performing experiments. M.A.F. wrote the majority of the text and performed the ilastik training. R.S. contributed portions of the text. P.S.M.M. contributed the pre-ilastik script. H.M. initially developed the MATLAB program, with refinement and final application by P.R. I.K. and G.G. supervised the research and contributed to writing the manuscript and drafting the figures. All authors edited the manuscript.

