## Supplementary material for "A Comprehensive Workflow for Imaging Live Insulin Secretion Events and Granules in Intact Islets": FIJI MACROS: Multi Point Plotter.ijm

```
// ImageJ Macro to plot points from a list of comma-separated xy coordinates and add them to ROI Manager

// Prompt user for xy coordinates
inputString = getString("Enter comma-separated xy coordinates:", "100,100,200,200,300,300");

// Split the input string and create arrays for x and y coordinates
coordinatesArray = split(inputString, ",");
xpoints = newArray(coordinatesArray.length / 2);
ypoints = newArray(coordinatesArray.length / 2);

// Populate xpoints and ypoints arrays
for (i = 0; i < coordinatesArray.length; i += 2) {
    xpoints[i / 2] = parseFloat(coordinatesArray[i]);
    ypoints[i / 2] = parseFloat(coordinatesArray[i + 1]);
}

// Create a new blank image with a white background
newImage("Point Plot", "32-bit White", 512, 512, 1);

// Create an empty ROI Manager
roiManager("reset");

// Loop through each point and add it as a single point ROI
for (i = 0; i < xpoints.length; i++) {
    // Create a single point selection
    makePoint(xpoints[i], ypoints[i]);

    // Add the single point selection to the ROI Manager
    roiManager("Add");
}

// Update the display
updateDisplay();
```
