## Supplementary material for "A Comprehensive Workflow for Imaging Live Insulin Secretion Events and Granules in Intact Islets": FIJI MACROS: Post ilastik processing.ijm

```
// Define the input directory and file

inputPath = "/Volumes/Ohi-Kaverina 4136-Data/Kaverina_Users/Jessica/coding_project/data/event/";
inputFile = "processed_Objects.h5.tif";
outputFolder = inputPath;  // Save results in the same folder

// import the .h5 file using Ilastik fiji plug in
run("Import HDF5", "select=/Users/myatpm/Desktop/irina/eventdata/processed_Objects.h5 datasetname=/exported_data axisorder=tzyxc");
selectImage("/Users/myatpm/Desktop/irina/eventdata/processed_Objects.h5/exported_data");

// convert image stack using look up table to glasbey

run("glasbey");

// copy
run("Duplicate...", "duplicate");

waitForUser;

setThreshold(0, 0);

// convert to binary
run("Make Binary", "background=Default calculate black create");

// analyze, set scale, remove scale, to make sure it is in pixels. 
// set measurements to include centroid 
// analyze, analyze particles, exclude edges, showe overlay
Image.removeScale;
run("Set Measurements...", "area centroid redirect=None decimal=3");
run("Analyze Particles...", "display exclude overlay add stack");


// open ROI manager
if (roiManager("count") == 0) {
    exit("No ROIs found in the ROI Manager.");
}

// clean up
run("Clear Results");

// loop through all the ROIs
for (i = 0; i < roiManager("count"); i++) {
    roiManager("Select", i); // Select ROI
    
    // get ROI name
    roiName = RoiManager.getName(i);
    
    // get data from string
    splitName = split(roiName, "-");
    
    // get frame number, first item
    frameNumber = splitName[0]; 

    // centroid
    run("Measure");

    // get centroid coordinates from the last added row in the Results Table
    rowIndex = nResults - 1;
    x = getResult("X", rowIndex);
    y = getResult("Y", rowIndex);
    
    // store frame number and coordinates in the Results Table
    setResult("Frame", rowIndex, frameNumber);
}

// show results
updateResults();
print("Macro completed successfully.");
```
