## Supplementary material for "A Comprehensive Workflow for Imaging Live Insulin Secretion Events and Granules in Intact Islets": FIJI MACROS: Pre ilastik processing.ijm

```
// Define the input directory and file
inputPath = "/Volumes/Ohi-Kaverina 4136-Data/Kaverina_Users/Jessica/coding_project/data/event/";
inputFile = "M3_Control_dmso_pseudo_islet_2_HG.nd2_C0.tif";
outputFolder = inputPath;  // Save results in the same folder

// Construct full input file path
path = inputPath + inputFile;

// open .tif
open(path);

// how many slices?
totalSlices = nSlices;

// auto adjust brightness and contrast
run("Enhance Contrast", "saturated=0.35");

// 1. create duplicate (from frame 2 to last)
firstRange = "2-" + totalSlices;
run("Duplicate...", "title=first.tif duplicate range=" + firstRange);

// 2. create second duplicate (from frame 1 to second-last)
secondRange = "1-" + (totalSlices - 1);
selectImage(inputFile);
run("Duplicate...", "title=second.tif duplicate range=" + secondRange);

// divide (first / second)
selectImage("first.tif");
imageCalculator("Divide create 32-bit stack", "first.tif", "second.tif");

// rename before saving
selectImage("Result of first.tif");
rename("processed");

// convert to grays
run("Grays");

// define output paths
processedTifPath = outputFolder + "processed.tif";
processedH5Path = outputFolder + "processed.h5";

// save .tif of processed
saveAs("Tiff", processedTifPath);

// export to .h5 using Ilastik 
run("Export HDF5", "output=[" + processedH5Path + "]");

run("Close All");
```
