## Supplementary material for "A Comprehensive Workflow for Imaging Live Insulin Secretion Events and Granules in Intact Islets": MATLAB Secretion clustering program: MATLAB Secretion Clustering Program.m

```
%This script was edited by Piilani Noguchi to assist Maggie Fye in the
%visualization of insulin granule secretion events. 5-28-2022


%The purpose of this script is to count the number of events that occur within set
%Region of Interest (ROI) boundaries as well as detecting clustering among
%those events.
%Inputed into the script are ROI coordinates in the form of csv files as
%well as coordinates of events, also in the form of a csv file.
%Outputed from the script is a figure displaying the ROIs as well as the
%events with clusters color-coded. Also outputed is an excel sheet with
%information about the ROI counts and clusters.


%These portions are manually written in. It is important that you are
%typing variables in properly.
Experiment_name = 'MATLAB Analysis'; %This is what you want to name your experiment this can be anything
Results_name = 'Flashes.xlsx'; %This is the name of the excel file with the flash data dont for get to put .filetype it this must be exact
Results_set_number = 'Movie 1'; %This is the name of the particular experiment (usually name of experiment trial usually a date or trial number) this can be anything
x_column = 2; %In the excel file with the flash data there is an X position column this is simply the number of that column (if in first column then x_column=1 and so on)
y_column = 3; %Similar as above... %In the excel file with the flash data there is an Y position column this is simply the number of that column (if in second column then x_column=2 and so on)
slice_column =4; % Similar as above (if Slice column is in third column then slice_column=3)


clearvars -except Experiment_name Results_name Results_set_number x_column y_column slice_column;
clf;
crop = '- Full Duration';
%% 

%Export Names
Export_name = append(Experiment_name,' - ', Results_set_number,' - Full Duration -Exported Data');
Excel_name = append(Export_name,'.xlsx');
Figure_name = append(Export_name,'.tiff');
%% 
%Create validation that x_column/y_column/slice_column are correctly being
%input?


%Read this results file
Results = readtable(Results_name); 
Results = Results{:,:};
x_coord = 6.25*Results(:,x_column); y_coord = 6.25*Results(:,y_column); time = Results(:,slice_column);
XY = [x_coord y_coord]; %Gives 2 column matrix of result coordinates
No_points = size(XY,1); %Gives number of points in the results


%% 

%Reading the ROI Coordinates
fnr = sprintf('*csv');
Files = dir(fnr);
ROI_number = length(Files);
%Files(length(Files))=[]; %Deleting the results .csv from the ROI files
%% 

%Matrix for ROI_counts
ROI1_counts2_nonclustered3_clustered4_numclusters5 = zeros(ROI_number,5);
for i = 1:ROI_number
    ROI1_counts2_nonclustered3_clustered4_numclusters5(i,1)=i;
end

%This matrix will contain all information about each individual result
%point
T1_x2_y3_roi4_posc5 = [time x_coord y_coord zeros(No_points,1)];

%This finds clusters in the result points based on x and y coordinates
%dbscan(X,epsilon,minpts)
if isempty(Results) ~= 1
    idx_position = dbscan(XY, 9, 3);
    idx_position(idx_position==-1)=0; %Changing -1 to 0 for non-cluster points
    %Input these index labels into the point matrix
    T1_x2_y3_roi4_posc5 = [T1_x2_y3_roi4_posc5 idx_position];
    No_clusters = max(idx_position); %This gives the number of clusters
end

if isempty(Results) == 1
    No_clusters = 0;
end

%Creates a matrix with the sizes of each cluster
Cluster1_size2 = zeros(No_clusters,2);
if isempty(Results) ~= 1
    for i=1:No_clusters
        Cluster1_size2(i,1)=i;
        A = find(T1_x2_y3_roi4_posc5(:,5)==i);
        Cluster1_size2(i,2)=size(A,1);
    end
end
%Round point coordaintes to an integer to check if they are contained in
%ROI coordinates
Points_rounded = round(XY);
%Reading each ROI pixel coordiantes in a loop
loc = zeros(No_points,1);

for fileno=1:ROI_number
    ROI_file_name = append(('ROI'),num2str(fileno),'.csv');
    ROI = readtable(ROI_file_name); ROI = ROI{:,:};
    X_ROI = ROI(:,1); Y_ROI = ROI(:,2);
    k=boundary(X_ROI, Y_ROI,1); plot(X_ROI(k),Y_ROI(k),'black')
    ROI_coord = [X_ROI Y_ROI];
    %Plot number of ROI on the Figure
    meanx = mean(X_ROI); meany = mean(Y_ROI);
    text(meanx,meany,num2str(fileno));
    %Detecting if the event resides in the ROI
    C = ismember(Points_rounded,ROI_coord,'rows');
    intersection_points=sum(C); ROI1_counts2_nonclustered3_clustered4_numclusters5(fileno,2) = intersection_points;
    loc(C)=fileno;
    hold on
end

ROI = csvread(Files(1).name,1,0);
x = ROI(:,1); y = ROI(:,2); k=boundary(x,y,1); plot(x(k),y(k),'black'); ROI=[x y];
meanx = mean(x);meany=mean(y);
Roi_number = string(1);
text(meanx,meany,Roi_number);

title(append(Experiment_name,' ',Results_set_number,crop));
T1_x2_y3_roi4_posc5(:,4)=loc.';
%% 

%Sort rows based on clusters so the legend is in order
if isempty(Results) ~= 1
    T1_x2_y3_roi4_posc5 = sortrows(T1_x2_y3_roi4_posc5,5);
    %Naming clusters for the figure legend
    a = T1_x2_y3_roi4_posc5(:,5);
    a = num2cell(a);
    a(cell2mat(cellfun(@(elem) elem == 0, a(:, :), 'UniformOutput', false))) = {'No Clstr'};
    a = string(a);
    
    %Plotting the points based on cluster label (for color coordination)
    fig = gscatter(T1_x2_y3_roi4_posc5(:,2),T1_x2_y3_roi4_posc5(:,3),a(:,1));
    Non_cluster = fig(1); Non_cluster.Color = 'k'; %makes points black if not in a cluster
    legend('Location','eastoutside');
    
end


axis equal;

hold off
%% 

%Find number of clustered and non-clustered events in each ROI
for i=1:ROI_number
    idx = T1_x2_y3_roi4_posc5(:,4)==i;
    a = T1_x2_y3_roi4_posc5(idx,:);
    if sum(idx) ~= 0
        ROI1_counts2_nonclustered3_clustered4_numclusters5(i,4)= sum(a(:,5)==0);
        ROI1_counts2_nonclustered3_clustered4_numclusters5(i,3)=sum(a(:,5)~=0);
    end
end
if isempty(Results)~=1
    %Finding the number of unique clusters
    Clusters = unique(T1_x2_y3_roi4_posc5(:,5)); Clusters = Clusters(Clusters ~= 0);
    No_clusters = size(Clusters,1);
    %Creates a matrix with the sizes and centers of each cluster
    Cluster1_sizes2_center34 = zeros(No_clusters,4);
    for i=1:No_clusters
        Cluster1_sizes2_center34(i,1) = i;
        A = find(T1_x2_y3_roi4_posc5(:,5)==i);
        Cluster1_sizes2_center34(i,2)=size(A,1);
    end
    z = zeros(No_clusters,2);
    for m = 1:No_clusters
        for i = 1:No_points
            %Creating a column vector of each of the points in cluster 'm'
            idx = T1_x2_y3_roi4_posc5(:,5)==m;
            w = T1_x2_y3_roi4_posc5(idx,:);
            %Column vectors of the x and y coordinates of the points in the
            %clusters
            x = w(:,2); y = w(:,3);
            meanx = mean(x); meany = mean(y);
            Cluster1_sizes2_center34(m,3)=meanx;
            Cluster1_sizes2_center34(m,4)=meany;
            Cluster_number = string(m);
            %Adds label to each of the clusters on the figure
        end
    end
    %Finding which ROI the cluster occurs in
    locC = zeros(No_clusters,1);
    Cluster_location_rounded = [round(Cluster1_sizes2_center34(:,3)) round(Cluster1_sizes2_center34(:,4))];
end
if isempty(Results)~= 1
    for fileno=1:ROI_number
        ROI_file_name = append('ROI',num2str(fileno),'.csv');
        ROI = csvread(ROI_file_name,1,0);
        x = ROI(:,1); y = ROI(:,2); ROI=[x y];
        C = ismember(Cluster_location_rounded,ROI,'rows'); intersection_points = sum(C);
        locC(C) = fileno;
        ROI1_counts2_nonclustered3_clustered4_numclusters5(fileno,5)= intersection_points;
        hold on
    end
end

set(findall(gcf,'-property','FontSize'),'FontSize',8);
% Turns the legend off if lots of clusters
if No_clusters > 25
    d = gca; legend(d,'off');
end
%% 

%Writing exported excel file
matrix1 = ROI1_counts2_nonclustered3_clustered4_numclusters5;
table1 = array2table(matrix1);
header1 = {'ROI Number','Total Counts','Cluster Counts','Non-Cluster Counts','Number of Clusters'};
cell1 = table2cell(table1);
cell1 = [header1; table2cell(table1)];
writecell(cell1,Excel_name,'Sheet',1);

matrix2 = T1_x2_y3_roi4_posc5;
matrix2 = sortrows(matrix2,1);
table2 = array2table(matrix2);
header2 = {'Time','Centroid X','Centroid Y','ROI','Cluster'};
cell2 = table2cell(table2);
cell2 = [header2; table2cell(table2)];
writecell(cell2,Excel_name,'Sheet',2);

matrix3 = Cluster1_size2;
table3 = array2table(matrix3);
header3 = {'Cluster Number','Counts'};
cell3 = table2cell(table3);
cell3 = [header3; table2cell(table3)];
writecell(cell3,Excel_name,'Sheet',3);


saveas(gcf,Figure_name, 'tiff');
```
